# Selection for translational efficiency in genes associated with alphaproteobacterial gene transfer agents

**DOI:** 10.1101/2022.06.17.496587

**Authors:** Roman Kogay, Olga Zhaxybayeva

## Abstract

Gene transfer agents (GTAs) are virus-like elements that are encoded by some bacterial and archaeal genomes. The production of GTAs can be induced by the carbon depletion and results in host lysis and release of virus-like particles that contain mostly random fragments of the host DNA. The remaining members of a GTA-producing population act as GTA recipients by producing proteins needed for the GTA-mediated DNA acquisition. Here, we detect a codon usage bias towards codons with more readily available tRNAs in the RcGTA-like GTA genes of alphaproteobacterial genomes. Such bias likely improves the translational efficacy during GTA gene expression. While the strength of codon usage bias fluctuates substantially among individual GTA genes and across taxonomic groups, it is especially pronounced in *Sphingomonadales*, whose members are known to inhabit nutrient-depleted environments. By screening genomes for gene families with similar trends in codon usage biases to those in GTA genes, we found a gene that likely encodes head completion protein in some GTAs were it appeared missing, and 13 genes previously not implicated in GTA lifecycle. The latter genes are involved in various molecular processes, including the homologous recombination and transport of scarce organic matter. Our findings provide insights into the role of selection for translational efficiency in evolution of GTA genes, and outline genes that are potentially involved in the previously hypothesized integration of GTA-delivered DNA into the host genome.

**Importance:** Horizontal gene transfer (HGT) is a fundamental process that drives evolution of microorganisms. HGT can result a rapid dissemination of beneficial genes within and among microbial communities, and can be achieved via multiple mechanisms. One peculiar HGT mechanism involves viruses “domesticated” by some bacteria and archaea (their hosts). These so-called gene transfer agents (GTAs) are encoded in hosts’ genomes, produced under starvation conditions, and cannot propagate themselves as viruses. We show that GTA genes are under selection to improve efficiency of their translation when the host activates GTA production. The selection is especially pronounced in bacteria that occupy nutrient-depleted environments. Intriguingly, several genes involved in DNA incorporation into a genome are under similar selection pressure, suggesting that they may facilitate integration of GTA-delivered DNA into the host genome. Our findings underscore the potential importance of GTAs as a mechanism of HGT under nutrient-limited conditions, which are widespread in microbial habitats.

## Introduction

Gene transfer agents are phage-like particles produced by multiple groups of bacteria and archaea (1). Unlike viruses, GTA particles tend to package random pieces of the host cell DNA instead of genes that encode GTAs themselves (2, 3). Released GTA particles can deliver the packaged genetic material to other cells (4), impacting exchange of genetic material in prokaryotic populations (5-7). The benefits of GTA production and GTA-mediated DNA acquisition are not well understood. It has been hypothesized that GTAs may facilitate DNA repair (8), enable population-level exchange of traits needed under the conditions of a nutritional stress via horizontal gene transfer (HGT) (5) or decrease population density during the carbon starvation periods (9).

To date, at least three independently exapted GTAs are functionally characterized (10). The most studied GTA system (RcGTA) belongs to the alphaproteobacterium *Rhodobacter capsulatus* (2). RcGTA is encoded by at least 24 genes that are distributed across 5 distinct genomic loci (11, 12). Seventeen of the 24 genes are situated in one locus, which is dubbed the ‘head-tail’ cluster because it encodes most of the structural proteins of the RcGTA particles (1). RcGTA-like ‘head-tail’ clusters are present in many alphaproteobacterial genomes; they evolve slowly and are inferred to be inherited mostly vertically from a common ancestor of an alphaproteobacterial clade that spans multiple taxonomic orders (12-14). Additionally, multiple cellular genes regulate RcGTA production, release and reception (11, 15). It is likely that other, yet undiscovered, genes in *R. capsulatus* genome are involved in GTA lifecycle.

Expression of RcGTA is known to be triggered by nutrient depletion (16), under which a small fraction of the *R. capsulatus* population becomes dedicated to GTA production (17, 18). As a result, RcGTA-producing cells likely express GTA genes at high levels. By extension, RcGTA-like GTA genes in other alphaproteobacteria (hereafter referred to as “GTA genes” for brevity) also likely to be highly expressed in GTA-producing cells of alphaproteobacterial populations.

Highly expressed genes that are involved in core biological processes, such as translational machinery, are known to exhibit a strong codon usage bias (19). For example, codon usage in ribosomal proteins, which are highly expressed in almost all organisms, deviates most dramatically from the distribution of codons expected under their equal usage corrected for organismal GC content (20). Such bias is primarily due to selection to match the pool of most abundant tRNA molecules in order to have the most efficient translation for proteins needed in high number of copies (21-23). As a result, highly expressed genes tend to have codons that correspond to the most abundant tRNA molecules in the cell. This type of selection is known as the “selection for translational efficiency” and is ubiquitous among bacteria (24).

Besides constitutively highly expressed genes, selection for translational efficiency also acts on genes that are highly expressed under specific environmental conditions that microorganisms experience (19, 24, 25). For instance, genes that utilize galactose have higher codon usage biases in budding yeasts that live in dairy-associated habitats than in yeasts that occupy alcohol-associated habitats (25). Additionally, genes that encode interacting proteins and genes involved in the same pathway often exhibit similar codon usage biases (25, 26).

In earlier work, we have discovered that alphaproteobacterial GTA genes have a striking bias towards GC-rich codons in comparison to the rest of the genome (9, 12). However, this bias is different from the codon usage bias: skewed composition of the encoded proteins towards containing energetically cheaper amino acids caused the first two positions of the codons to be enriched in Gs and Cs, due to the structure of the genetic code (9). In this study, we examined GTA genes in 208 alphaproteobacterial genomes and assessed if there is additionally a codon *usage* bias due to the genes being under selection for the translation efficiency. For this purpose, we used two well-established metrics for assessment of codon usage bias and its match to the tRNA abundance: effective number of codons (ENC) (20) and tRNA adaptation index (tAI) (27).

ENC quantifies how equally synonymous codons are used in a gene, and varies from 20 (when only one codon is used per each amino acid; strong bias) to 61 (when all codons are used equally; no bias). The tAI measures how optimally the codon usage of each gene fits the available tRNA pool by correlating frequency of each codon in the gene with the abundance of its cognate tRNA. The degree of adaptation of a gene is gauged by comparing its tAI value to the tAI values of all other genes in a genome. We also searched for genes whose involvement in GTA production and regulation is currently unsuspected by screening GTA-encoding genomes for genes with codon usage patterns similar to those of GTA genes.

## Results

### Codon usage bias of GTA genes and its match to available tRNAs varies across GTA genes and GTA-containing genomes

To assess the presence of codon usage bias in GTA genes across alphaproteobacteria, we have calculated ENC for each “reference GTA gene” (see **Methods** for the definition) and compared them against the expected ENC of a gene in a genome under the null model of no codon usage bias, corrected for the genomic GC content (28). Indeed, we found that 1,543 out of 2,308 (66.8%) reference GTA genes detected across 208 GTA head-tail clusters deviate from the genome-specific null expectations by more than 10% (**Supplemental Figure S1**). However, there is a substantial variation in this deviation for different GTA genes (**Supplemental Figure S2**), and only in genes *g5* and *g8* the deviation is significantly higher than the genomic average (Kruskal-Wallis rank sum test, p-value < 2.2e-16; Dunn’s test, p-value < 0.05, Benjamini-Hochberg correction).

To assess the match of the observed codon usage bias to available tRNA pool, we calculated tAI values of the reference GTA genes across 208 genomes and converted them to percentile tAI values (ptAI; see **Methods** for the definition) to allow for the intergenomic comparisons. Similar to the ENC values, the ptAI values also vary substantially across the genes and genomes (**Figure 1**), suggesting that the strength of selection for translational efficiency should be examined in individual GTA genes and in specific taxonomic groups, which we investigate in the next two sections.

**Figure 1.**
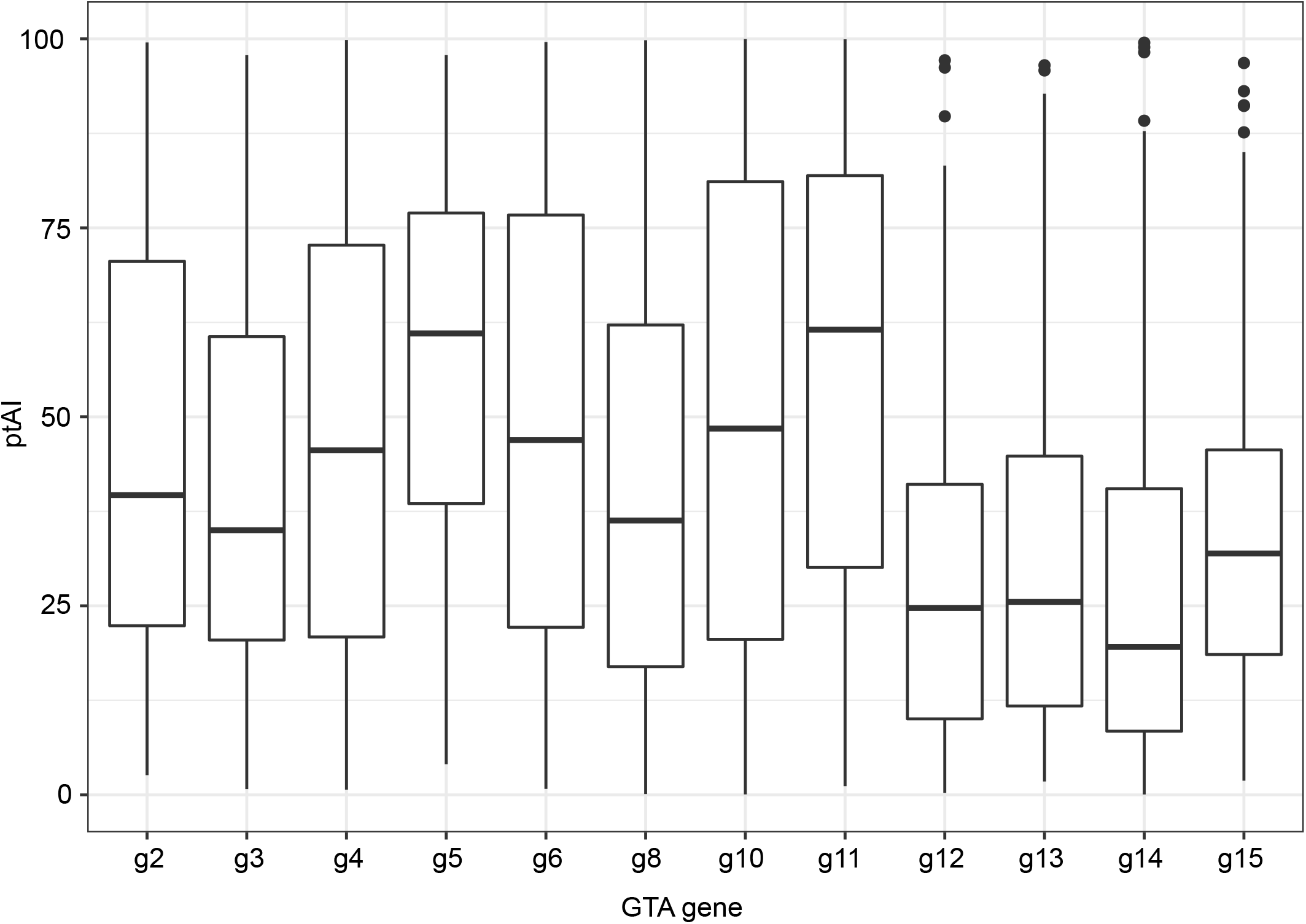
Distribution of ptAI values among reference GTA genes from GTA head-tail clusters in 208 alphaproteobacterial genomes. Line within a box displays the median ptAI value for a GTA gene across all genomes, in which the gene was detected. The boxes are bounded by first and third quartiles. Whiskers represent ptAI values within 1.5*interquartile range. Dots outside of whiskers are outliers.

### Selection for translational efficiency is uneven among GTA genes

The differences of ptAI values among the reference GTA genes are statistically significant (Kruskal-Wallis rank sum test, p-value < 2.2e-16) (**Figure 1**). Particularly notable is a significant decline in ptAI values of the region encoding genes *g12* through *g15* (Dunn’s test, p-value < 0.05, Benjamini-Hochberg correction), which are located at the 3’ end of the head-tail cluster and encode the tail components of a GTAs particle. In contrast, ptAI values of the genes *g5* (encoding major capsid protein) and *g11* (encoding tail tape measure protein) are significantly higher than ptAI values of other GTA genes (Dunn’s test, p-values < 0.05, Benjamini-Hochberg correction). Notably, protein g5 is detected in the largest number of copies (145) per RcGTA particle than any other protein (29), while proteins g12-g15 are present in a small number of copies (1-6) per RcGTA particle (29). Given that genes encoding proteins needed in a larger number of copies have a higher degree of adaptation to the tRNA pool (30), we hypothesize that the observed variation in ptAI values of GTA genes reflects the different number of GTA proteins in a GTA particle. Protein g11, however, is found in only 3 copies per RcGTA particle (29) and therefore a demand for a larger copy number cannot explain its high ptAI values.

Variation of ptAI values could also be due to physical location of the genes in the GTA head-tail cluster. Similar to the operons (31), genes in the RcGTA head-tail cluster are co-transcribed from a single promoter upstream of the cluster (15, 32). Because genes at the 3’ end of operons tend to have lower expression levels (33), the low ptAI values of GTA genes *g12-g15* may be due to their distant location from the promoter.

### Selection for translational efficiency is the strongest in *Sphingomonadales*’ genomes

In addition to variability in ptAI values across different GTA genes, ptAI values of individual GTA genes vary substantially across the 208 genomes (**Figure 1**). To evaluate if these differences represent variation in selection pressure in distinct taxonomic groups, we initially examined the ptAI values of gene *g5* that were grouped by alphaproteobacterial order. The *g5* gene was chosen due to its high abundance of the encoded protein in RcGTA particles (more copies than all other structural proteins combined) and for being the only gene with the highest detected deviations from the average genomic values for both ENC and ptAI. We found that ptAI values of the *g5* gene vary significantly among members of the four alphaproteobacterial orders (Kruskal-Wallis rank sum test, p-value < 0.05) (**Figure 2A**). In particular, *g5* genes from the *Sphingomonadales*’ genomes have significantly higher ptAI values than those from genomes of bacteria from other three orders (Mann-Whitney U test, p-value < 0.05, Benjamini-Hochberg correction). Twelve of the fourteen *g5* genes with the highest overall ptAI values (> 90) (**Figure 2A**) also belong to the *Sphingomonadales* genomes. Beyond just *g5* gene, all reference GTA genes, as a group, have higher ptAI values in *Sphingomonadales* than in members of the three other alphaproteobacterial orders (Mann-Whitney U test, p-value < 0.05, Benjamini-Hochberg correction) (**Supplemental Figure S3**). These observations suggests that in *Sphingomonadales* in particular, there is a strong selection for efficient production of GTA particles. Because *Sphingomonadales* are known to live in nutrient-depleted environments (34), we suggest that GTA production is especially beneficial in those habitats to exert strong selection for translational efficiency.

**Figure 2.**
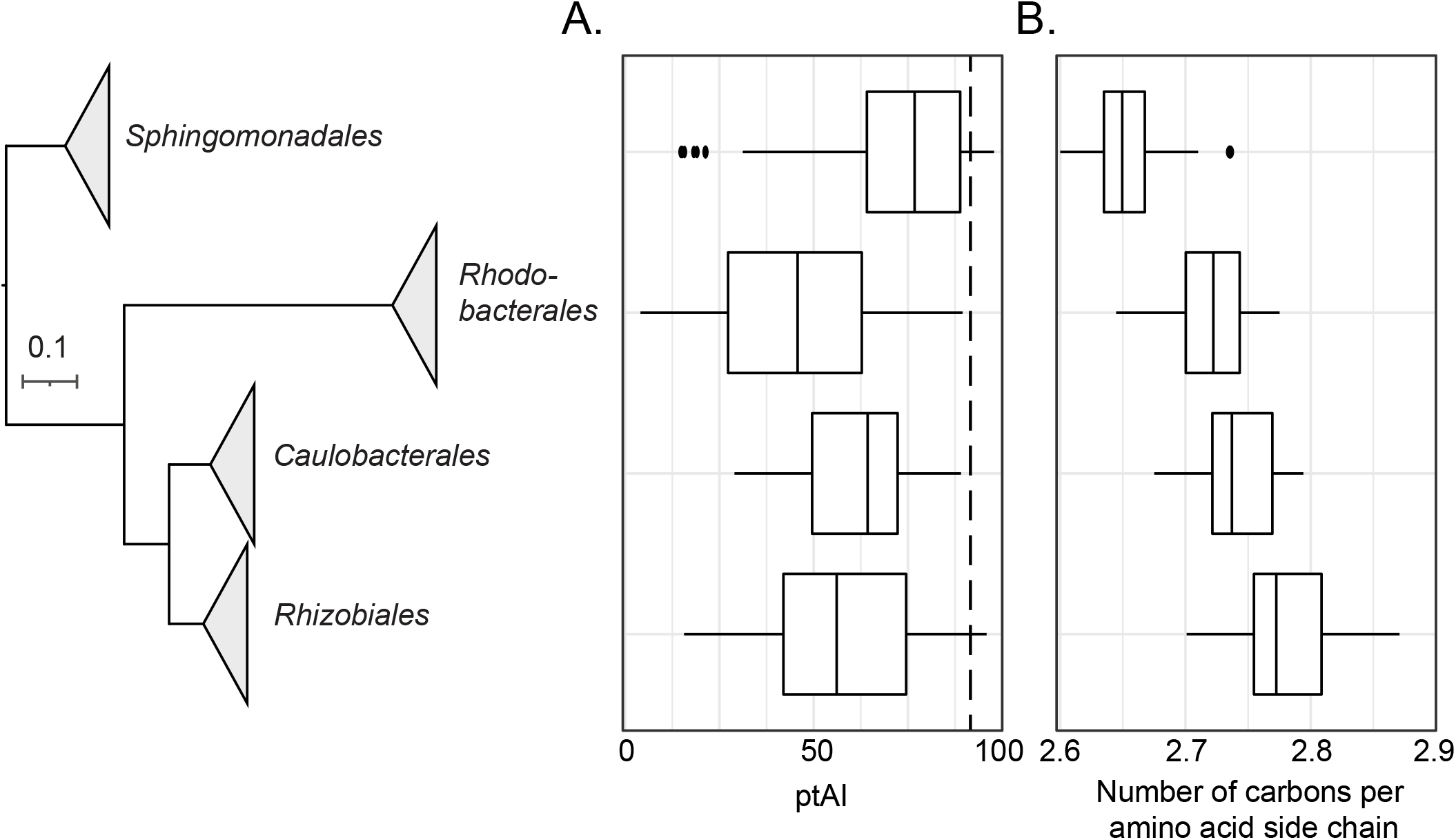
Distributions of (A) ptAI values in the major capsid protein-encoding gene (*g5*) and (B) carbon content of amino acids in the g5 protein across four orders of the class *Alphaproteobacteria*. On both panels A and B, line within a box displays the median ptAI value for g5 representatives within an order. The boxes are bounded by first and third quartiles. Whiskers represent ptAI values within 1.5*interquartile range. Dots outside of whiskers are outliers. The phylogenetic tree on the Y-axis is the reference phylogenomic tree (see **Methods** for details), in which branches are collapsed at the taxonomic order level. Dashed line in Panel A marks ptAI value of 90.

### The increase in translational efficiency of GTA genes is associated with a reduced energetic cost for production of the encoded proteins

Among the GTA proteins in four alphaproteobacterial orders, *Sphingomonadales’* GTA proteins also have the strongest skew in amino acid composition towards energetically cheaper amino acids (**Figure 2B**). To evaluate if selection for energy efficiency is linked to selection for translational efficiency, we examined the relationship between the ptAI values of GTA genes and the number of carbons in amino acid chains encoded by the *Sphingomonadales* GTA genes. We found that there is a significant negative correlation between them (Pearson R = -0.19, N = 636, p-value < 0.05). We propose that in *Sphingomonadales* benefits associated with production of GTA particles in nutrient-limited conditions led not only to the selection for translational efficiency, but also to the selection for use of energetically cheaper amino acids in the GTA genes.

### Fourteen gene families have translational efficiency trends similar to those of GTA genes

While the strength of selection for translational efficiency acting on GTA genes varies across gene and genomes, we found that the combinations of ptAI values across all reference GTA genes in a genome have similar trends across the 208 genomes. The similarity is significant in all pairwise reference GTA gene comparisons (**Supplemental Figure S4)**, as determined using phylogenetic generalized least squares (PGLS) method. We conjecture that ptAI values of genes in the other loci of a GTA “genome”, as well as the host genes involved in GTA lifecycle, would exhibit similar trends to the ptAI values of reference GTA genes, allowing for discovery of yet unsuspected genes involved in GTA lifecycle. To identify such unknown genes that may be co-expressed with GTA genes, we examined correlations of ptAI values between reference GTA genes and 3,477 other gene families present in 208 alphaproteobacterial genomes. The PGLS analysis revealed 14 gene families, whose ptAI values correlate significantly with ptAI values of the reference GTA genes (**Table 1)**.

**Table 1.** Functional annotations of 14 gene families, whose ptAI values have a significantly similar trend to ptAI values of the reference GTA genes.

| Gene | RefSeq ID of a Representative Protein | RefSeq Record Annotation | COG Category | COG functional category description |
| --- | --- | --- | --- | --- |
| *gafA* | WP_121690074.1 | DUF6456-domain containing protein | K | Transcription |
| *addA* | WP_121690807.1 | Double-strand break repair helicase AddA | L | Replication, recombination, and repair |
| *addB* | WP_121690808.1 | Double-strand break repair protein AddB | L | Replication, recombination, and repair |
| *xseA* | WP_092770100.1 | Exodeoxyribonuclease VII large subunit | L | Replication, recombination, and repair |
| *dinG* | WP_067681212.1 | ATP-dependent DNA helicase | KL | Transcription; Replication, recombination, and repair |
| *hrpB* | WP_121690814.1 | ATP-dependent helicase HrpB | L | Replication, recombination, and repair |
| *priA* | WP_121691035.1 | Primosomal protein N' | L | Replication, recombination, and repair |
| *glnE* | WP_121690099.1 | Bifunctional [glutamine synthetase] adenylyltransferase/[glutamine synthetase]-adenylyl-L-tyrosine phosphorylase | OT | Molecular chaperones and related functions; Signal transduction mechanism |
| *ccmE* | WP_010971299.1 | Cytochrome c maturation protein CcmE | O | Molecular chaperones and related functions |
| *ATP12* | WP_092769070.1 | ATP12 family chaperone protein | O | Molecular chaperones and related functions |
| *tonB* | WP_119082607.1 | Energy transducer TonB | M | Cell wall/membrane/envelope biogenesis |
| *TPR* | WP_162687979.1 | Tetratricopeptide repeat protein | M | Cell wall/membrane/envelope biogenesis |
| *smrA* | WP_010970599.1 | Smr/MutS family protein | S | Function unknown |
| *crtB* | WP_121690324.1 | Phytoene/Squalene synthase family protein | I | Lipid transport and metabolism |

One of 14 identified gene families is a homolog of *gafA*, which encodes a crucial transcription activator of GTA particles production in *Rhodobacter capsulatus* (11, 15). This gene is located outside of the RcGTA’s head-tail cluster, and therefore was not included in the set of reference GTA genes, but its discovery demonstrates the suitability of our approach to identify genes linked to the GTA lifecycle. Interestingly, *gafA* homologs were previously described only in the genomes of *Rhodobacterales* and some *Rhizobiales* (11, 12, 15). However, with different criteria in the OrthoFinder-based similarity searches, we were able to identify this regulator in 196 of the 208 genomes (94.7%), spanning all GTA-containing alphaproteobacterial orders. The evolutionary history of the *gafA* homologs is largely congruent with the reference phylogenomic tree (normalized quartet score of 0.87) and even more so with the phylogeny of the concatenated GTA reference genes (normalized quartet score of 0.92) (tree topologies are available at https://doi.org/10.6084/m9.figshare.20082749), suggesting that the *gafA* gene had co-evolved with the GTA ‘head-tail’ cluster since the last common ancestor of RcGTA-like GTAs.

The remaining 13 gene families belong to several functional categories of the Clusters of Orthologous Groups (COG) classification (**Table 1**). While proteins encoded by some of these genes can be postulated to be involved in GTA lifecycle (exemplified below by the *addAB, xseA*, and *tonB* genes), similarity of codon usage biases between other genes and reference GTA genes can be explained by their expression at similar environmental conditions (exemplified by three genes from ‘molecular chaperones and related functions’ COG category).

Protein products of the *addA* and *addB* genes form the heterodimeric helicase-nuclease complex that repairs double-stranded DNA breaks by homologous recombination and is functionally equivalent to the RecBCD complex (35). The knockout of the AddAB complex is associated with a deficiency in RecA-dependent homologous recombination (36). We hypothesize that the *addAB* pathway is involved in recombination of GTAs’ genetic material with the host’s genome.

The main function of exodeoxyribonuclease VII large subunit (xseA), which is encoded by the *xseA* gene, is to form a complex with xseB and degrade single-stranded DNA to oligonucleotides. However, expression of *xseA* gene without *xseB* gene leads to cell death (37). Because we did not find any correlation of codon usage bias between the *xseB* gene and GTA genes, we speculate that instead of involvement in processing of GTA DNA, *xseA* gene product facilitates lysis of GTA producing cells and release of GTA particles.

The *tonB* gene encodes tonB energy transducer. TonB-dependent transporters are involved in transport of diverse compounds, including carbohydrates, amino acids, lipids, vitamins and iron (38-40). Similar to the quorum-sensing regulated expression of the gene encoding GTA receptor in the non-GTA-producing cells of a *Rhodobacter capsulatus* population (4), the *tonB* gene could also be regulated to be expressed in the non-GTA-producing cells to aid the uptake of the nutrients released from the lysed cells via TonB-dependent transporters. The *tonB* gene is currently detected only in members of *Sphingomonadales* order, suggesting that such nutrient uptake is most relevant in the nutrient-limited environments.

Three genes from the ‘molecular chaperones and related functions’ COG category are less likely to be directly involved in GTA lifecycle, because GTAs already encode their own chaperones that assist GTA protein folding (29). However, it is well known that chaperones tend to be highly expressed in bacteria at times of stress and facilitate the survival of cells in rapidly changing environmental conditions (41). Because chaperones are essential in responding to the starvation-induced cellular stresses (42), we conjecture that observed similarity in ptAI values of the reference GTA genes is due to their expression being triggered by the similar environmental conditions.

To evaluate if the detected gene families interact with each other and with GTA genes, we have constructed the protein-protein interaction network of the 14 gene families, 12 GTA reference genes and 50 additional interactor proteins from the STRING database (**Figure 3**). Thirteen of the 14 families and all 12 reference GTA genes belong to two protein-protein interaction sub-networks (**Figure 3**), one of which contains all GTA reference genes, while the other is involved in a wide range of functions (**Table 1**). By carrying out the KEGG enrichment analysis, we found significant overrepresentation of four molecular pathways in the second protein-protein interaction network (**Supplemental Table S1**). Consistent with the 6 of the 13 gene families being assigned to the “replication, recombination, and repair” COG category, two of the KEGG pathways are ‘homologous recombination’ and ‘mismatch repair’, further corroborating involvement of identified genes in integration of the genetic material delivered by GTAs into recipients’ genomes. Two additional pathways, ‘carotenoid biosynthesis’ and ‘terpenoid backbone biosynthesis’, are less likely to be directly involved in the lifecycle of GTAs. Production of secondary metabolites is known to be protective against stress factors (43, 44), and carbon starvation leads to the upregulation of carotenoid biosynthesis pathway (45, 46). Similar to the above-described genes encoding chaperones, we hypothesize that expression of ‘carotenoid biosynthesis’ and ‘terpenoid backbone biosynthesis’ genes is not related to GTA lifecycle, but is initiated by conditions that also activate production of GTAs.

**Figure 3.**
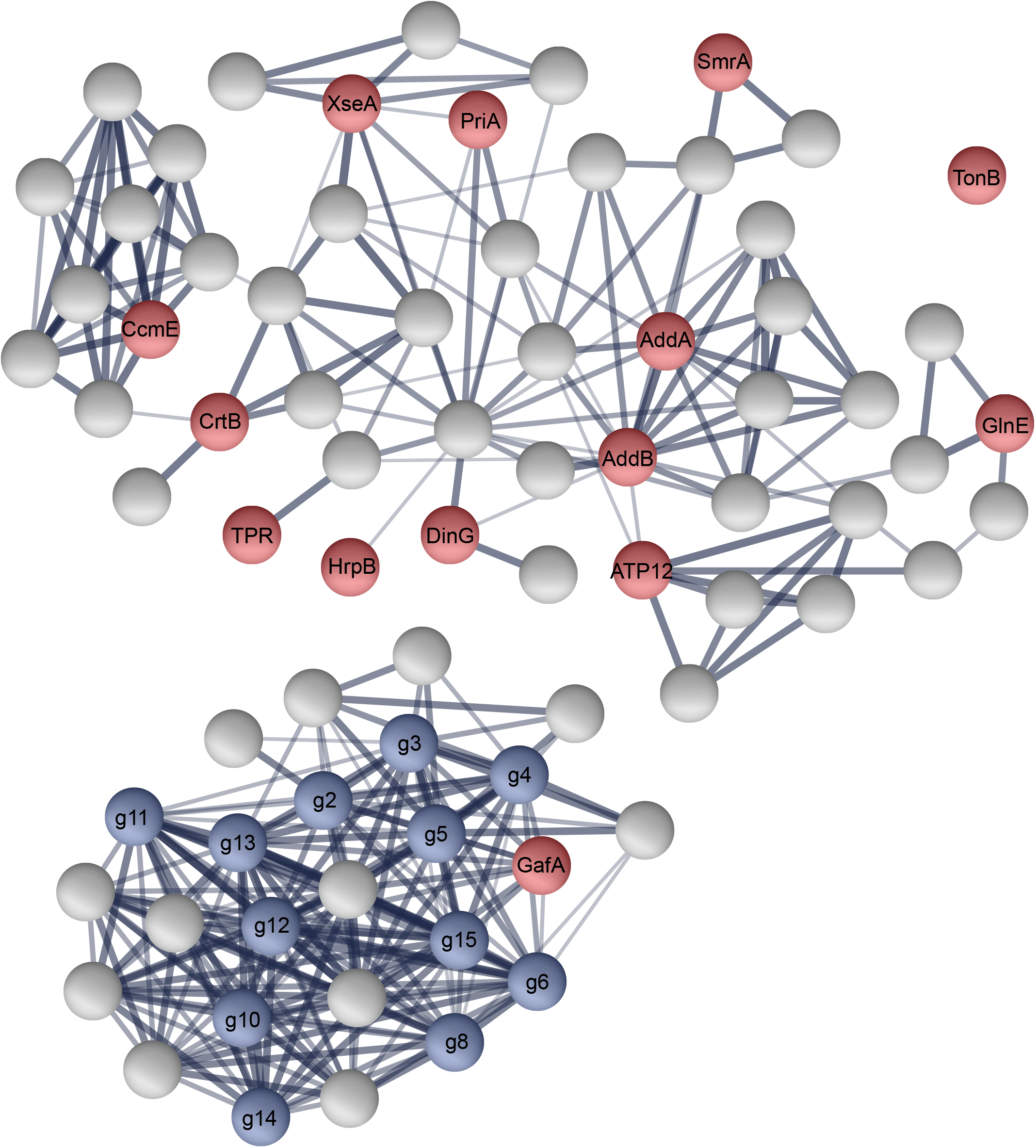
The protein-protein interactions among 12 GTA reference proteins and 14 proteins putatively co-expressed with GTAs. Nodes represent individual proteins. Blue-colored nodes correspond to GTA reference proteins and red-colored nodes correspond to 14 putatively co-expressed proteins. Gray-colored nodes represent additional proteins found through the STRING functional enrichment analysis. The thickness of the edges is proportional to the STRING’s confidence score of protein interactions (varying between 0.4 [thin line] and 1.0 [thick line]).

### A replacement of the head completion protein in *Sphingomonadales*’ GTAs

Gene content of GTA head-tail clusters varies across alphaproteobacteria (12). While some clusters do not contain homologs of all RcGTA genes, others include additional genes that are conserved across multiple clusters but have no known function (12, 13). To predict whether any of these additional genes play a role in GTA production, we compared their ptAI values of genes found in at least 10 genomes to ptAI values of the reference GTA genes. One gene family, which is found only within GTA head-tail clusters of 11 genomes in one subclade of *Sphingomonadales* (GenBank accessions are available at https://doi.org/10.6084/m9.figshare.20082749), has a significant positive correlation with 5 out of the 12 GTA reference genes (**Supplemental Table S2**). Interestingly, within *Sphingomonadales* GTA head-tail clusters this gene is located where the *g7* gene, which encodes a head completion protein, is found in the RcGTA head-tail cluster (**Supplemental Figure S5**). Only seven of the 55 *Sphingomonadales* genomes in our dataset have detectable homologs of the *g7* gene. Among the remaining 48 genomes, 22 contain a gene encoding a protein of unknown function in the “gene *g7* locus”, while 26 genomes don’t have any gene in that locus.

The members of the identified gene family are substantially shorter than the RcGTA gene *g7* and have a different secondary structure (**Figure 4**), precluding the possibility that the identified protein is simply too divergent for a detectable amino acid similarity. However, we found viral head completion proteins that have similar protein length and similar secondary structures to both GTA head completion protein and the identified gene family (**Figure 4**). We conjecture that the gene encoding the head completion protein was replaced in some *Sphingomonadales* by a gene encoding an analogous viral protein.

**Figure 4.**
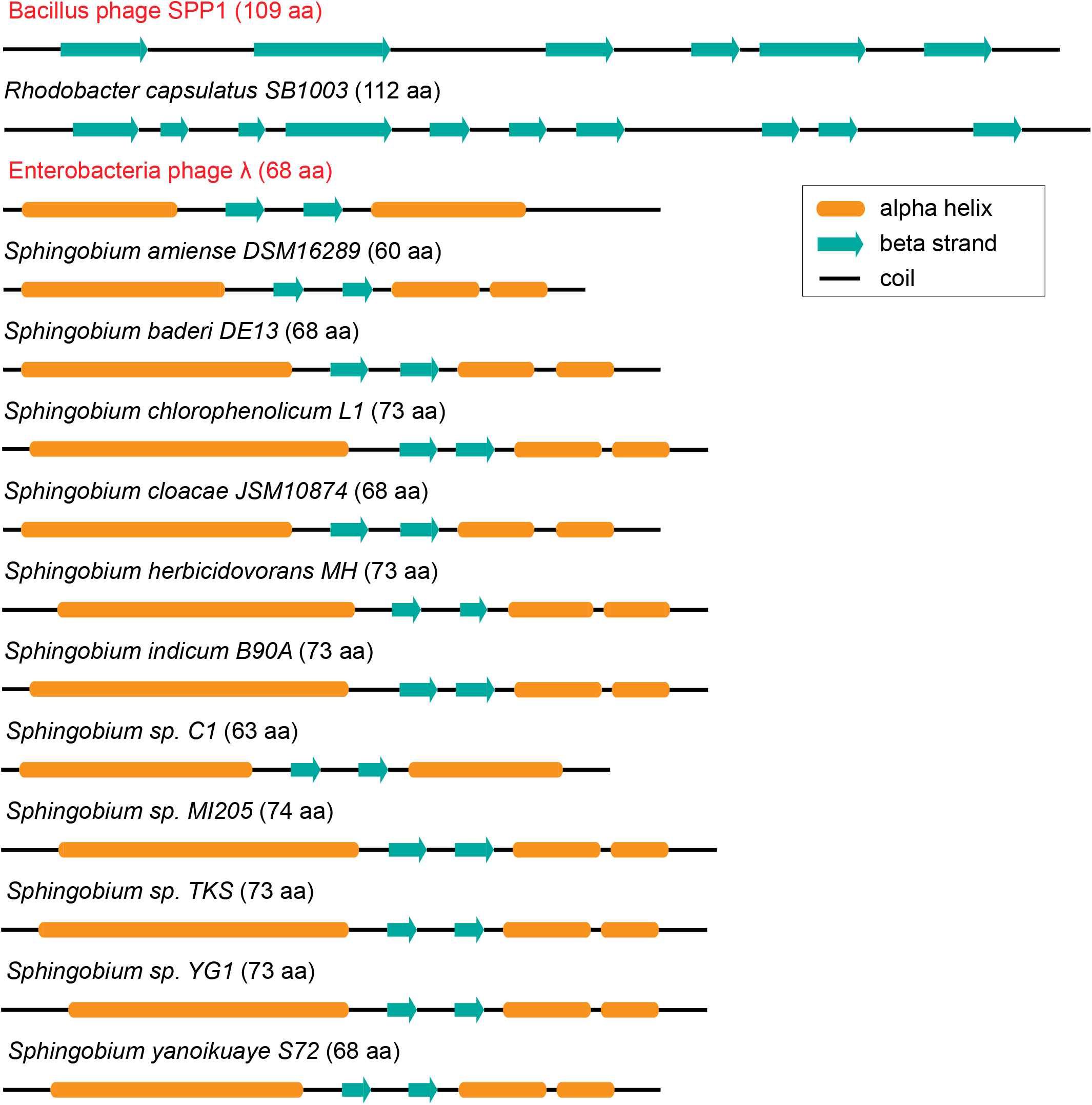
Secondary structures of head completion proteins from phages and GTAs. The Enterobacteria phage lambda gpW and Bacillus phage SPP1 (highlighted in red) are two representatives of viral head completion proteins with major differences in lengths and secondary structures. The secondary structures of *R. capsulatus* (PDB ID 6TUI_8), Enterobacteria phage lambda gpW (1HYW), and Bacillus phage SPP1 (2KCA) proteins were retrieved from the PDB database. The secondary structures of the putative head completion proteins from *Sphingomonadales* were predicted computationally. The secondary structures are scaled with respect to the protein lengths, which are listed in parentheses next to the taxonomic names.

## Discussion

Our analyses of codon usage biases suggest that alphaproteobacterial GTA systems are under selection for an optimal translation of GTA proteins from GTA genes achieved by using codons with more readily available tRNAs. The strength of such selection for translational efficiency is the most pronounced (and therefore most easily detectable) in the major capsid protein gene, which is needed to be expressed to produce thousands of copies per GTA-producing bacterium. Additionally, the strength of the selection for translational efficiency varies across taxonomic groups, but is particularly prominent in *Sphingomonadales* order, whose members typically inhabit nutrient-limited conditions. We hypothesize that the observed variation in the selection strength depends on severity and duration of the nutrient scarcity experienced by a population capable of producing GTAs. On the one hand, a long-term exposure to nutrient-depleted conditions would trigger a more efficient and/or more frequent production of GTA particles, which would lead to a greater survival of communities with a better translational efficiency of GTA systems and thus a higher codon usage bias in the GTA genes. On the other hand, if GTAs are needed only in rare occasions due to the stable and abundant nutrient supplies, the selection for translational efficiency would be weak and would result in a lower codon usage bias. Combined with an observation that production of GTAs is triggered by the nutritional stress (16), our findings that the selection is the strongest in alphaproteobacteria that inhabit nutrient-limited environments further underscore the earlier hypothesized importance of GTA systems in situations of nutrient scarcity (9).

Additionally, the stronger selection for translation efficiency in GTA genes is associated with a larger decline in the carbon content of the proteins the genes encode. These findings suggest that benefits associated with GTA production are substantial enough to drive selection for both translational efficiency and low energetic costs of the translated proteins. We speculate that these modifications of GTA proteins allow the bacterial population under adverse conditions to increase both the speed of GTA particle production and the number of released GTA particles.

We hypothesized that genes that are located outside of the GTA head-tail cluster, but are involved in GTA lifecycle, including processing and integration of the GTA-delivered DNA, would have signatures of selection for translational efficiency similar to those of GTA genes.

Gratifyingly, our genome-wide screen for such patterns detected the direct GTA activator gene, *gafA* (15). We also identified multiple genes not yet implicated in GTA lifecycle. Several of these genes are involved in recombination and mismatch repair, providing bioinformatic evidence for the hypothesis that GTAs facilitate HGT by distributing genetic fragments that become incorporated into recipients’ genomes via homologous recombination (4). Involvement of other genes with similar selection pressures in GTA lifecycle is speculative and needs to be investigated experimentally. But the putative co-expression of *xseA* and *tonB* genes with GTA genes raises an intriguing possibility that, in addition to HGT, GTA production may provide an extra benefit in a nutrient-depleted environment: scavenging of scarce organic matter from GTA-producing cells. The lysis of the GTA-producing cells could be mediated by XseA and their debris could be imported as nutrients by the surviving cells via TonB-dependent transporters.

Alphaproteobacterial GTAs likely originated millions of years ago from a lysogenic phage, and since then they were mostly vertically inherited by many alphaproteobacterial lineages (12, 14). However, similar to the HGT influence onto many other regions of a typical bacterial genome (47), it is very likely that over time GTAs experienced gene replacements via HGT (12). Instances of HGT between GTAs and phages have been already documented (11, 48). By examining the patterns of selection for translational efficiency, we identified another case of likely ancient gene exchange with viruses that resulted in the replacement of the gene encoding head completion protein in some *Sphingomonadales*. Curiously, the gene currently has no significant primary sequence similarity to any gene in GenBank. Many other unannotated ORFs in alphaproteobacterial head-tail clusters outside of *Rhodobacterales* (12) may also have functional roles in their respective GTA regions. Notably, when alphaproteobacterial RcGTA-like genomic regions appear incomplete due to lack of many homologs to genes required for GTA production in *R. capsulatus*, it could be due our inability to recognize some genes due to their replacements with analogous genes. Because such incomplete RcGTA-like clusters are abundant in alphaprotebacteria (12), GTAs could be morphologically diverse and even more widespread across alphaproteobacteria than we currently estimate (13).

## Methods

### Dataset of representative alphaproteobacterial genomes with GTA head-tail clusters

As an initial data set, we selected 212 representative alphaproteobacterial genomes previously predicted to contain GTAs (9). The gene annotations of the genomes were downloaded from the RefSeq database (49) in October 2020. GTA head-tail clusters (1) were predicted using the GTA-Hunter program (13). Because GTA-Hunter identifies only 11 out of the 17 genes in the RcGTA’s head-tail cluster and also requires genes to align with their RcGTA homologs by at least 60% of their length, some GTA genes were likely missed by GTA-Hunter. To look for these potential false negatives, additional BLASTP (50) searches with the e-value cutoff of 0.1 were performed using 17 RcGTA head-tail cluster genes as queries and protein-coding genes in 212 genomes as a database. Only matches located within the genomic regions designated as GTA gene clusters by GTA-Hunter were kept. In four genomes, calculations of genes’ adaptation to tRNA pool (see **“Evaluation of the adaptiveness of protein-coding genes to the tRNA pool”** section below for details) did not converge. As a result, only 208 genomes were retained in the reported analyses (GenBank accessions are available at https://doi.org/10.6084/m9.figshare.20082749).

### Identification of gene families in 208 alphaproteobacterial genomes

Within each genome, protein-coding genes less than 300 nucleotides in length were excluded in order to reduce the stochasticity of codon usage bias values due to the insufficient number of codons. The remaining protein-coding genes were clustered into gene families using Orthofinder v2.4 (51) with default parameters and DIAMOND (52) for the amino acid sequence similarity search. Only gene families detected in at least 40 genomes were retained to ensure statistical power.

Some alphaproteobacterial GTA head-tail cluster regions contain protein-coding ORFs that do not have significant similarity to the RcGTA homologs of the genes shown to be required for GTA production in RcGTA. Gene families of these ORFs were retrieved from the collection of gene families predicted for all protein-coding genes (regardless of their length) using Orthofinder v2.4 (51) with default parameters and DIAMOND (52) for the amino acid sequence similarity search. Only gene families that are both located within the genomic region encoding GTA head-tail cluster and found in at least 10 genomes were retained.

### Reference set of GTA genes

Although RcGTA head-tail cluster contains 17 genes, genes *g3*.*5* and *g10*.*1* are less than 300 nucleotides in length, and genes *g1* and *g7* are not detected widely across analyzed genomes. Additionally, codon usage patterns of gene *g9* were found to be very different from other GTA genes (see **“Examination of similarity in adaptation to the tRNA pool among GTA genes”** section below for details). Therefore, in our inferences about selection, we considered only 12 of the 17 GTA genes (**Supplemental Table S3**), which we designate throughout the manuscript as “GTA reference genes”.

Amino acid sequences of GTA reference genes were aligned individually using MAFFT-linsi v7.455 (53) and then concatenated into a single alignment. Each gene was treated as a separate partition in the alignment and the best substitution model for each gene was determined by ModelFinder (54). The maximum likelihood tree was reconstructed using IQ-TREE v1.6.7 (55) and the support values were calculated via 1,000 ultrafast bootstrap replicates (56).

### Reconstruction of the reference phylogenomic tree

Twenty-nine marker proteins that are present in a single copy in more than 95% of the 208 retained genomes were retrieved using AMPHORA2 (57). Amino acid sequences within each of the 29 marker families were aligned using MAFFT-linsi v7.455 (53). The best substitution matrix for each family was determined by *ProteinModelSelection*.*pl* script downloaded from https://github.com/stamatak/standard-RAxML/tree/master/usefulScripts in October 2020. Individual alignments of the marker families were concatenated, but each alignment was treated as a separate partition with its own best substitution model in the subsequent phylogenetic reconstruction. The maximum likelihood tree was reconstructed using IQ-TREE v1.6.7 (55) and the support values were calculated using 1,000 ultrafast bootstrap replicates (56).

### Evaluation of codon usage bias in protein-coding genes using “effective number of codons” metric

For the retained genes in each genome, effective number of codons (ENC) (20) and G+C content variation at the 3rd codon position in the synonymous sites (GC3s) were calculated using CodonW (http://codonw.sourceforge.net). The null model of no codon usage bias was calculated as described in dos Reis et al. (28) using an in-house script (available in the FigShare repository; see below). For every gene, the deviation of its ENC from the null model was calculated using the in-house script. Genes that have observed ENC higher than the expected were excluded from analyses.

### Evaluation of the adaptiveness of protein-coding genes to the tRNA pool

The tRNA genes in each genome were predicted using tRNAscan-SE v 2.06, using a model trained on bacterial genomes (58, 59) and the Infernal mode without HMM filter to improve the sensitivity of the search (60). tRNA gene copy number was used as the proxy for tRNA abundance, following the previously reported observation that the two correlate strongly (28, 61). The adaptiveness of each codon (ω_i_) to the tRNA pool was calculated using the stAIcalc program with the maximum hill climbing stringency (62). The tRNA adaptation index (tAI) of each retained gene was calculated as the geometric mean of its ω_i_ values (27). Because the distribution of tAI values varies among genomes (63) (**Supplemental Figure S6**), tAI values were converted to their relative percentile tRNA adaptation index (ptAI) within a genome. The ptAI values range between 0 and 100, and represent the percentage of analyzed genes in a genome that have a smaller tAI than a particular gene.

### Examination of similarity in adaptation to the tRNA pool among GTA genes

The ptAI values were retrieved for a subset of 13 GTA genes that are at least 300 nucleotide in length and are widely detected across all taxonomic groups. The linear regression analysis of ptAI values between all GTA gene pairs was conducted using the phylogenetic generalized least squares method (PGLS) (64). The reference phylogenomic tree was used to correct for the shared evolutionary history. The analysis was done using the ‘caper’ package (65) and λ, δ and κ parameters were estimated using the maximum likelihood function. Because ptAI values of gene *g9* were not significantly correlated with the ptAI values of 8 out of the 12 other examined GTA genes at p-value cutoff of 0.001 (**Supplemental Table S4**), the gene *g9* was not included into the reference set of GTA genes.

### Identification of genes with ptAI values similar to that of the GTA genes

For each gene family, the “within-genome” ptAI values were retrieved. For gene families with at least two paralogs, the ptAI values for all paralogs from a particular genome were replaced with their median ptAI value.

To identify gene families that exhibit tRNA pool adaptation patterns similar to those of GTA genes, a linear regression model of ptAI values between these gene families and reference GTA genes was fit using the PGLS (64). The PGLS analysis was carried out using the ‘caper’ package (65) and λ, δ and κ parameters were estimated using the maximum likelihood function. The reference phylogenomic tree was used to correct for the shared phylogenetic history. For gene families found in at least 40 genomes, a gene family was designated to be associated with a GTA, if obtained fit of the model was statistically significant across all reference GTA genes. Because gene families found in less than 40 genomes were kept only if the genes are located within the genomic regions encoding GTA head-tail clusters (see the “Identification of gene families in 208 alphaproteobacterial genomes” section), a more relaxed criterion was adopted for such gene families: a gene family was designated to be associated with a GTA if the fit of the model was statistically significant across at least 40% of reference GTA genes. If a significantly associated gene family contained paralogs, the PGLS analysis was repeated by using individual ptAI values across all possible combinations of paralogs (if the total number of combinations was < 1,000) or across random 1,000 combinations of paralogs (if the total number of combinations was > 1,000). This was carried out to ensure that the detected signal was not due to sampling associated with selecting the median ptAI value.

Genes with a significant similarity in trend of ptAI values were annotated via eggNOG-mapper v2.1 (66).

### Protein-protein interaction of GTA genes and gene families with similar ptAI values

To identify protein-protein interaction networks, reference GTA genes and genes from families with similar tRNA pool adaptation patterns were retrieved from the *Sphingomonas* sp. MM1 genome, chosen for it being the only genome that contains all genes from the GTA reference gene set and all 14 gene families listed in **Table 1**. The locus tags of the retrieved *Sphingomonas* sp. MM1 genes were used as queries against STRING database v 11.0b (last accessed July 2021) (67) with the medium confidence score cutoff and all active interaction sources. The retrieved protein-protein interaction network was visualized in STRING using the queries and up to 50 additional interactor proteins, and displaying edges based on the STRING confidence scores. The KEGG pathways (68) enrichment analysis was conducted via hypergeometric testing on the whole retrieved network, as implemented in STRING.

### Analysis of other protein-coding genes situated within GTA head-tail clusters

For gene families within GTA head-tail clusters, ptAI values were retrieved and compared to ptAI values of the reference GTA gene set using PGLS analysis as described above. For the only gene family with a significant association with GTA genes, the secondary structure of its proteins were predicted using Porter v5.0 (69). To retrieve available viral head completion proteins, the phrase ‘head-completion protein’ was used as a query against the UniProt database (accessed in August 2021) (70). Among the 24 manually annotated (“reviewed”) matches from the Swiss-Prot sub-database of the UniProt database, only 2 viral matches (accession numbers P68656 and P68660) had length similar to the genes in the above described gene family. Both proteins belong to the λ phage gpW family, and for *Escherichia* phage λ protein 3D structure is available in PDB (71). The secondary structure of RcGTA’s g7 protein, structural viral homolog of RcGTA’s g7 from *Bacillus* phage SPP1 (gp16) (29) and head completion protein of phage λ were retrieved from the PDB database (71) in August 2021.

In 48 *Sphingomonadales* genomes without a detectable homolog of RcGTA gene *g7*, the genomic space either between the homologs of the RcGTA genes *g6* and *g8*, or, in genomes without *g6* homolog, between homologs of the RcGTA genes *g5* and *g8*, was searched for presence of open reading frames.

### Refinement of the *tonB* gene family using phylogenetic tree

To identify orthologs within the large *tonB* gene family, evolutionary history of the *tonB* gene family was reconstructed and evaluated. To do so, amino acid sequences of the *tonB* gene family were aligned using MAFFT-linsi v7.455 (53). The phylogeny was reconstructed in IQ-TREE v1.6.7 (55) using the best substitution model (LG+F+R6) detected by ModelFinder (54). The tree was visualized using the iTOL v6 (72). The phylogeny was used to subdivide the family into two families, whereas the five genes on very long branches served as an outgroup (tree topology is available at https://doi.org/10.6084/m9.figshare.20082749).

### Calculation of energetic cost associated with production of the encoded proteins

To quantify the energetic cost of proteins, the carbon content of their amino acids was used as a proxy and was calculated by counting the number of carbons in the amino acid side chains, as described in Kogay et al. (9). The total number of carbons in each protein was normalized by the protein length.

### Retrieval and phylogenetic analyses of *gafA* homologs

Amino acid sequences of *gafA* homologs in 196 alphaproteobacterial genomes were detected via Orthofinder (gene family OG0001218). Only homologs found in single copy in a genome (194 in total) were retained for phylogenetic analysis. These homologs were aligned using MAFFT-linsi v7.455 (53). The best substitution model (LG+F+R6) was determined by ModelFinder (54) and the maximum-likelihood tree was reconstructed by IQ-TREE v1.6.7 (55) with the number of iterations to stop set to 500. The support values were calculated using 1,000 ultrafast bootstrap replicates (56). Both GTA reference tree and reference phylogenomic tree were pruned to match the taxa in *gafA* phylogeny. The normalized quartet scores were calculated using ASTRAL v5.7.8 (73).

### Data availability

The following data are available in the FigShare repository under DOI 10.6084/m9.figshare.20082749 (https://doi.org/10.6084/m9.figshare.20082749): accession numbers of 208 analyzed alphaproteobacterial genomes; accession numbers of the GTA regions identified in the analyzed genomes; accession numbers of genes in gene families across analyzed genomes; raw data related to tAI and ENC calculations; an in-house script for ENC calculations; slopes and p-values of associations detected in PGLS analyses; accession numbers of the putative g7 proteins in *Sphingomonadales* genomes; multiple sequence alignments and phylogenetic trees of *tonB* and *gafA* gene families, concatenated phylogenomic markers, and concatenated GTA reference genes.

## Acknowledgements

This work was supported in part by the Simons Foundation Investigators in Mathematical Modeling of Living Systems program (Award #327936 to O.Z.).

## Supplemental Material Captions

Supplemental Tables S1-S4 and Figures S1-S6 are provided as PDF files.

**Supplemental Table S1. Four molecular pathways significantly over-represented in the protein-protein interaction network shown in Figure 3**. The pathway information was obtained from KEGG database.

**Supplemental Table S2. Significance and slope of the fit of the phylogenetic generalized least squares (PGLS) models between the reference GTA genes and putative head-completion protein in *Sphingomonadales***. Statistically significant associations (p-value <0.05) are highlighted in orange.

**Supplemental Table S3. Decisions behind choosing the RcGTA ‘head-tail’ cluster homologs for the reference GTA gene set**. The GTA genes selected for the reference set are highlighted in orange. Functional annotations are based on the descriptions in the RefSeq database records, unless noted otherwise.

**Supplemental Table S4. Significance and slope of the fit of the phylogenetic generalized least squares (PGLS) models between the reference GTA genes and the *g9* gene**.

**Supplemental Figure S1. Distribution of deviations of the effective number of codon (ENC) values from the expected ENC values under the null model of no codon bias**. The distribution contains deviations for 2,308 reference GTA genes found in the 208 genomes. Numbers on the plot designate the number of reference GTA genes in an interval delineated by dashed lines.

**Supplemental Figure S2. Deviation of the effective number of codon (ENC) values for individual reference GTA genes in comparison to the genomic average**. The deviation of the ENC from the expectation under the null model for each GTA gene was normalized by the average ENC deviation of its genome. Line within a box displays the median normalized ENC value for a GTA gene across all genomes. The boxes are bounded by first and third quartiles. Whiskers represent ptAI values within 1.5*interquartile range. Dots outside of whiskers are outliers.

**Supplemental Figure S3. Distributions of ptAI values in all reference GTA genes across four orders of the class *Alphaproteobacteria***. Line within a box displays the median ptAI value for a GTA gene across all genomes. The boxes are bounded by first and third quartiles. Whiskers represent ptAI values within 1.5*interquartile range. Dots outside of whiskers are outliers.

**Supplemental Figure S4. PGLS model fit among ptAI values of the reference GTA gene pairs**. Each pairwise comparison is represented by a rectangle that is color-coded according to the p-values from the PGLS analysis of the reference GTA gene pairs. The numerical p-values are listed within each rectangle.

**Supplemental Figure S5. Gene neighborhood of the RcGTA gene *g7* and of the putative *g7* replacement gene in 11 *Sphingomonadales* spp**. Only region corresponding to the RcGTA ‘head-tail’ cluster is depicted, with each gene represented by an arrow scaled relative to its length within each cluster. The RcGTA genes (*g1*-*g15*) are color-coded and their homologs in *Sphingomonadales* are shown in the same color. Putative *g7* replacements in *Sphingomonadales* are shown in magenta and marked with an arrow. Pseudogenes are colored in black, while genes without an established relationship to GTA production are shown in gray. Phylogenetic tree is a subtree extracted from the reference phylogeny. The scale bar corresponds to the number of substitutions per site.

**Supplemental Figure S6. Distribution of tAI values in protein-coding genes of the analyzed genomes**. Only genes at least 300 nucleotides in length were included. **A**. Distribution of tAI values of genes in three representative alphaproteobacterial genomes, selected to have the lowest, the median, and the highest mean tAI value among 208 genomes. **B**. Distribution of the average genomic tAI values across 208 genomes.

