## Supplemental Tables and Figure for "Selection for translational efficiency in genes associated with alphaproteobacterial gene transfer agents"

**Supplemental Table S1. Four molecular pathways significantly over-represented in the protein-protein interaction network shown in Figure 3.** The pathway information was obtained from KEGG database.

| KEGG Pathway ID | Pathway name | p-value (after Benjamini-Hochberg correction) |
|---|---|---|
| sphm00900 | Terpenoid backbone biosynthesis | 0.0102 |
| sphm03440 | Homologous recombination | 0.0119 |
| sphm00906 | Carotenoid biosynthesis | 0.0202 |
| sphm03430 | Mismatch repair | 0.0349 |

**Supplemental Table S2. Significance and slope of the fit of the phylogenetic generalized least squares (PGLS) models between the reference GTA genes and putative head-completion protein in *Sphingomonadales*.** Statistically significant associations (p-value <0.05) are highlighted in orange.

| Reference GTA gene | p-value | Slope |
|---|---|---|
| *g2* | 0.82086 | -0.03579 |
| *g3* | 0.59683 | 0.08254 |
| *g4* | 0.40305 | 0.13635 |
| *g5* | 0.21417 | 0.15219 |
| *g6* | 0.01450 | 0.41291 |
| *g8* | 0.83146 | 0.03253 |
| *g10* | 0.50256 | 0.17411 |
| *g11* | 0.00004 | 0.10553 |
| *g12* | 0.03593 | 0.25124 |
| *g13* | 0.85847 | 0.02868 |
| *g14* | 0.00170 | 0.59603 |
| *g15* | 0.00002 | 0.13779 |

**Supplemental Table S3. Decisions behind choosing the RcGTA 'head-tail' cluster homologs for the reference GTA gene set**. The GTA genes selected for the reference set are highlighted in orange. Functional annotations are based on the descriptions in the RefSeq database records, unless noted otherwise.

| GTA gene | RcGTA RefSeq ID | RcGTA functional annotation | Included? | Reason for a gene exclusion |
|---|---|---|---|---|
| *g1* | WP_013067406.1 | small terminase [1] | | Homologs are detected only in *Rhodobacterales* order |
| *g2* | WP_031321187.1 | terminase family protein | Yes | |
| *g3* | WP_013067408.1 | phage portal protein | Yes | |
| *g3.5* | WP_031323538.1 | hypothetical protein | | The gene is < 300 nucleotides in length |
| *g4* | WP_013067410.1 | HK97 family phage prohead protease | Yes | |
| *g5* | WP_037091462.1 | phage major capsid protein | Yes | |
| *g6* | WP_013067412.1 | adaptor protein [2] | Yes | |
| *g7* | WP_013067413.1 | head-tail adaptor protein | | Detected only in 10% of *Sphingomonadales* genomes |
| *g8* | WP_013067414.1 | tail terminator protein [2] | Yes | |
| *g9* | WP_013067415.1 | phage major tail protein, TP901-1 family | | Patterns of the selection for translational efficiency are inconsistent with those of other GTA genes |
| *g10* | WP_013067416.1 | gene transfer agent family protein | Yes | |
| *g10.1* | WP_013067417.1 | phage tail assembly chaperone | | The gene is < 300 nucleotides in length |
| *g11* | WP_013067418.1 | phage tail tape measure protein | Yes | |
| *g12* | WP_013067419.1 | distal tail protein [2] | Yes | |
| *g13* | WP_013067420.1 | baseplate hub protein [2] | Yes | |
| *g14* | WP_013067421.1 | peptidase | Yes | |
| *g15* | WP_013067422.1 | glycoside hydrolase/phage tail family protein | Yes | |

1. Sherlock, D., J.X. Leong, and P.C.M. Fogg, *Identification of the First Gene Transfer Agent (GTA) Small Terminase in Rhodobacter capsulatus and Its Role in GTA Production and Packaging of DNA.* J Virol, 2019. **93**: e01328-19.
2. Bardy, P., et al., *Structure and mechanism of DNA delivery of a gene transfer agent.* Nat Commun, 2020. **11**: 3034.

**Supplemental Table S4. Significance and slope of the fit of the phylogenetic generalized least squares (PGLS) models between the reference GTA genes and the *g9* gene.**

| Reference GTA gene | p-value | Slope |
|---|---|---|
| *g2* | 0.01287 | 0.18307 |
| *g3* | 0.00145 | 0.20857 |
| *g4* | 0.00099 | 0.25720 |
| *g5* | 2.23E-06 | 0.28841 |
| *g6* | 0.00511 | 0.23485 |
| *g8* | 0.00645 | 0.20851 |
| *g10* | 0.12627 | 0.08297 |
| *g11* | 0.00061 | 0.21435 |
| *g12* | 0.00031 | 0.31015 |
| *g13* | 0.00162 | 0.26218 |
| *g14* | 0.03469 | 0.14122 |
| *g15* | 0.00138 | 0.27938 |

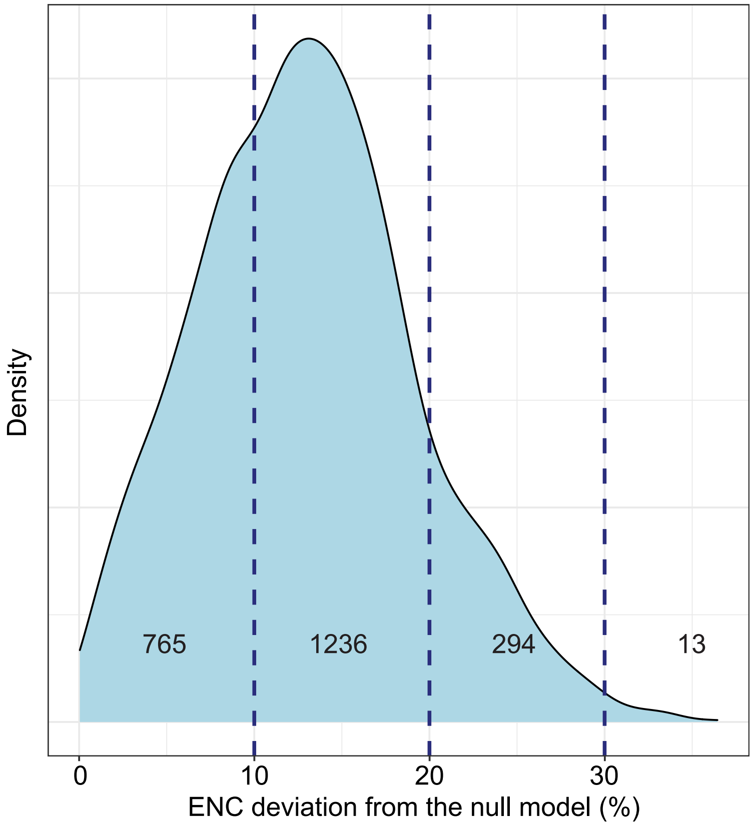


**Supplemental Figure S1. Distribution of deviations of the effective number of codon (ENC) values from the expected ENC values under the null model of no codon bias.** The distribution contains deviations for 2,308 reference GTA genes found in the 208 genomes. Numbers on the plot designate the number of reference GTA genes in an interval delineated by dashed lines.

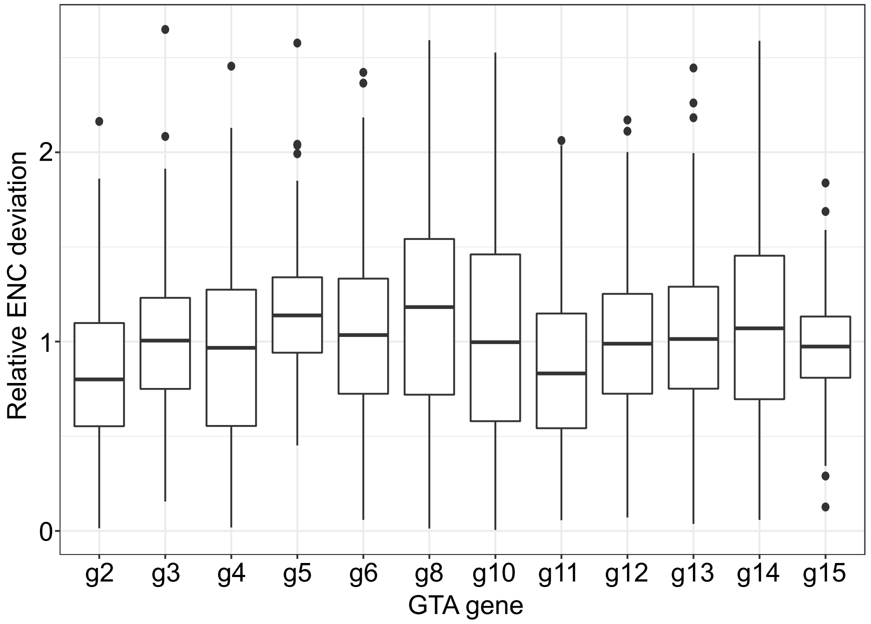


**Supplemental Figure S2. Deviation of the effective number of codon (ENC) values for individual reference GTA genes in comparison to the genomic average.** The deviation of the ENC from the expectation under the null model for each GTA gene was normalized by the average ENC deviation of its genome. Line within a box displays the median normalized ENC value for a GTA gene across all genomes. The boxes are bounded by first and third quartiles. Whiskers represent ptAI values within 1.5*interquartile range. Dots outside of whiskers are outliers.

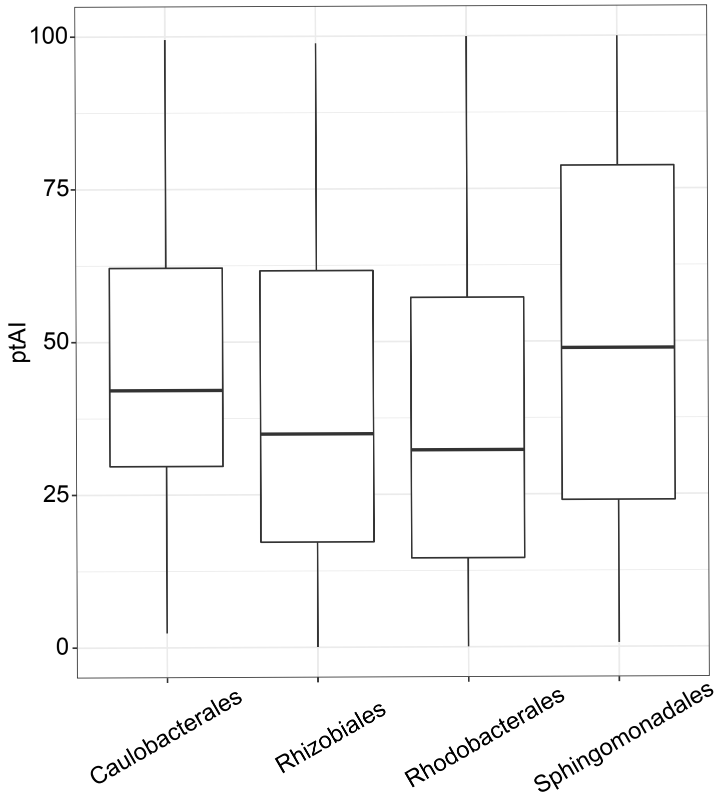


**Supplemental Figure S3. Distributions of ptAI values in all reference GTA genes across four orders of the class *Alphaproteobacteria*.** Line within a box displays the median ptAI value for a GTA gene across all genomes. The boxes are bounded by first and third quartiles. Whiskers represent ptAI values within 1.5*interquartile range. Dots outside of whiskers are outliers.

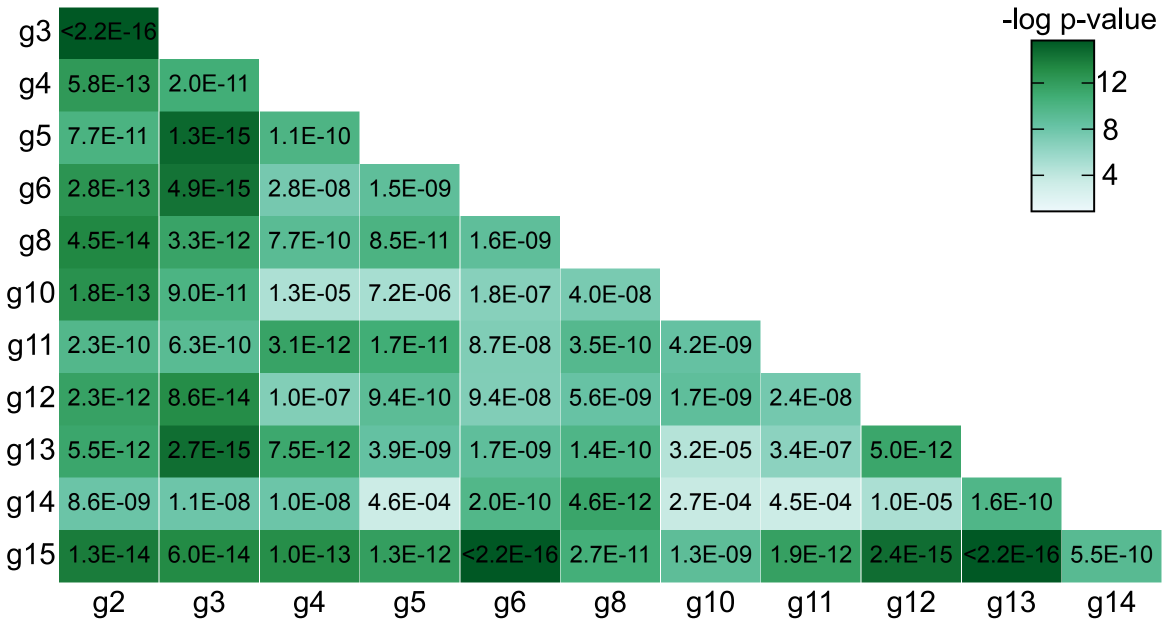


**Supplemental Figure S4. PGLS model fit among ptAI values of the reference GTA gene pairs.** Each pairwise comparison is represented by a rectangle that is color-coded according to the p-values from the PGLS analysis of the reference GTA gene pairs. The numerical p-values are listed within each rectangle.

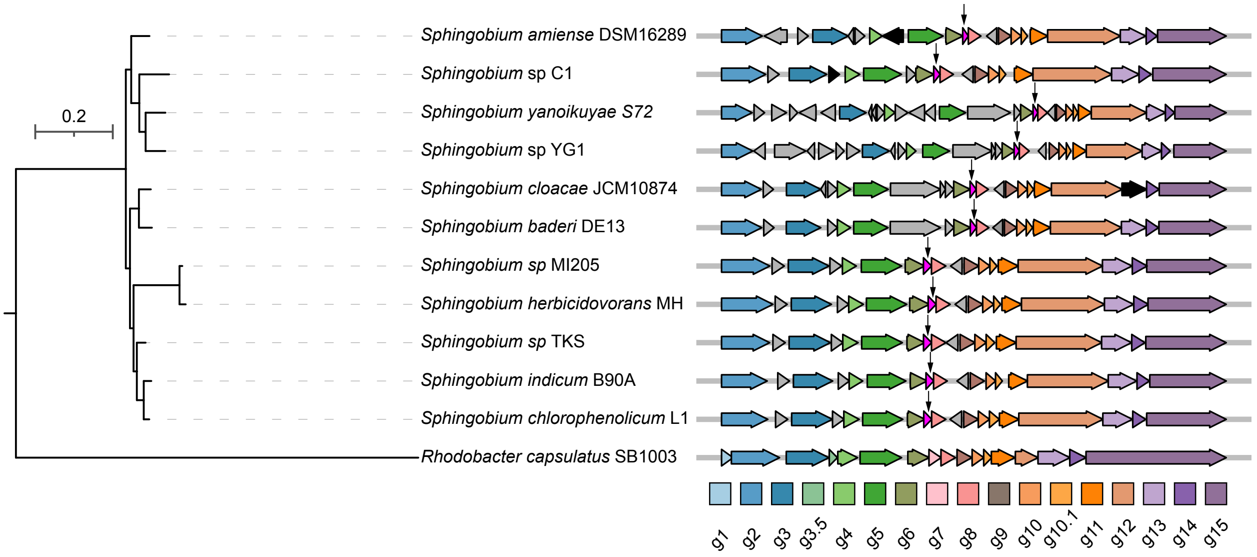


**Supplemental Figure S5. Gene neighborhood of the RcGTA gene *g7* and of the putative *g7* replacement gene in 11 *Sphingomonadales* spp.** Only region corresponding to the RcGTA ‘head-tail’ cluster is depicted, with each gene represented by an arrow scaled relative to its length within each cluster. The RcGTA genes (*g1*-*g15*) are color-coded and their homologs in *Sphingomonadales* are shown in the same color. Putative *g7* replacements in *Sphingomonadales* are shown in magenta and marked with an arrow. Pseudogenes are colored in black, while genes without an established relationship to GTA production are shown in gray. Phylogenetic tree is a subtree extracted from the reference phylogeny. The scale bar corresponds to the number of substitutions per site.

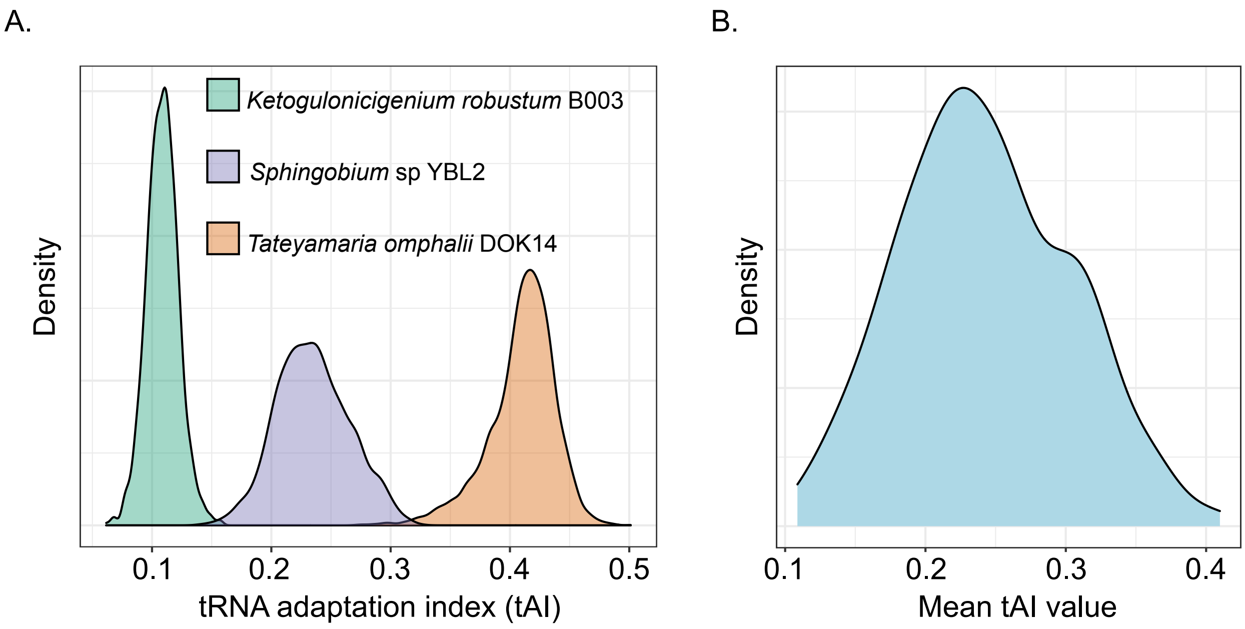


**Supplemental Figure S6. Distribution of tAI values in protein-coding genes of the analyzed genomes.** Only genes at least 300 nucleotides in length were included. **A.** Distribution of tAI values of genes in three representative alphaproteobacterial genomes, selected to have the lowest, the median, and the highest mean tAI value among 208 genomes. **B.** Distribution of the average genomic tAI values across 208 genomes.
